# Advanced transcriptomic analysis reveals the role of efflux pumps and media composition in antibiotic responses of *Pseudomonas aeruginosa*

**DOI:** 10.1101/2022.04.04.487074

**Authors:** Akanksha Rajput, Hannah Tsunemoto, Anand V. Sastry, Richard Szubin, Kevin Rychel, Siddharth M. Chauhan, Joe Pogliano, Bernhard O. Palsson

## Abstract

*Pseudomonas aeruginosa* is an opportunistic pathogen and major cause of hospital acquired infections. The pathogenicity and virulence of *P. aeruginosa* is largely determined by its transcriptional regulatory network (TRN). We used 411 transcription profiles of *P. aeruginosa* from diverse growth conditions to construct a quantitative TRN by identifying independently modulated sets of genes (called iModulons) and their condition-specific activity levels. The current study focused on the use of iModulons to analyze pathogenicity and antibiotic resistance of *P. aeruginosa*. Our analysis revealed: 1) 116 iModulons, 81 of which show strong association with known regulators; 2) novel roles of two-component systems in regulating antibiotics efflux pumps; 3) substrate-efflux pump associations; 4) differential iModulon activity in response to beta-lactam antibiotics in bacteriological and physiological media; 5) differential activation of ‘Cell Division’ iModulon resulting from exposure to different beta-lactam antibiotics; and 6) a role of the PprB iModulon in the stress-induced transition from planktonic to biofilm lifestyle. In light of these results, the construction of an iModulon-based TRN provides a transcriptional regulatory basis for key aspects of *P. aeruginosa* infection, such as antibiotic stress responses and biofilm formation. Taken together, our results offer a novel mechanistic understanding of *P. aeruginosa* pathogenicity.

**Significance:** Large data sets and machine learning are impacting a growing number of areas of research in the life sciences. Once the compendia of bacterial transcriptomes reached a critical size, we could use source signal extraction algorithms to find lists of co-regulated genes (called iModulons) associated with a transcription factor (TF) to them. The gene composition of iModulons and their condition-dependent activity levels constitute a quantitative description of the composition of bacterial transcriptomes. This study shows how this approach can be used to reveal the responses of *P. aeruginosa* to antibiotics and thus yield a deep regulatory understanding of pathogenicity properties. This study motivates the execution of similar studies for the other ESKAPEEs to yield a broad understanding of the role of TRNs in antibiotic responses to these urgent threat bacterial pathogens.

## Introduction

The transcriptional regulatory network (TRN) for any organism controls the expression of genes in response to genetic and environmental perturbations. For pathogenic bacteria, the TRN coordinates important functions for the infection of hosts, such as virulence mechanisms, antibiotic resistance, and biofilm formation ^1,2^. Uncovering the TRN of pathogenic bacteria can be helpful in identifying novel drug targets and the underlying molecular mechanisms of inhibition. *Pseudomonas aeruginosa* is an important member of the ESKAPEE group of pathogens that are notoriously multi-drug resistant ^3^. In addition to its intrinsic antibiotic resistance mechanisms ^3^, the large genome and genetic plasticity of *P. aeruginosa* ^4^ allows the bacteria to grow in varied environments but complicates comprehensive TRN mapping.

Previous efforts to characterize TRNs from *Pseudomonas* species are consolidated on the ‘Pseudomonas Genome DB,’ which annotated 371 confirmed or putative transcription factors (TFs) in the *P. aeruginosa* PAO1 genome ^5^. Efforts have been made to elucidate the TRNs mainly by the characterization of TFs using experimental approaches like transcriptional profiling, chromatin immunoprecipitation, and other similar methods ^6,7^. Further, Wang *et al* validated the TFs in *P. aeruginosa* involved in virulence using the high-throughput systematic evolution of ligands by exponential enrichment assay ^8^. These studies describe the mechanistic details about the TF binding sites and regulon annotation. However, these methods are expensive, time-consuming, and not derived from the transcriptional profiles. Although these studies are necessary for providing a foundation for understanding the TRN of *P. aeruginosa*, other approaches harness big data to generate a comprehensive understanding of the TRN. Therefore, in the current study, we use transcriptional profiles of *P. aeruginosa* to reconstruct the TRN.

Independent component analysis (ICA) is used to separate multivariate signals into additive components. For transcriptomic datasets, it identifies independently modulated sets of genes, called iModulons, and quantifies their activity under specific conditions. Since iModulons capture the signals from transcriptional regulators, they can be compared to regulons. Regulons are sets of co-regulated genes defined based on bottom-up approaches using a variety of biomolecular methods, whereas iModulons are defined in a top-down manner using machine learning of entire transcriptomic profiles. Previously, we used ICA to annotate the TRNs of *Escherichia coli* ^9^, *Staphylococcus aureus* ^10^, *Bacillus subtilis* ^11^, and *Sulfolobus acidocaldarius* ^12^, which generated valuable hypotheses including putative regulatory interactions, novel associations between regulators and the conditions which may activate them, and specific insights into transcriptomic reallocation during key physiological processes. ICA has also been used to study the effect of adaptive laboratory evolution on the TRN ^13,14^. Results from all public iModulon analyses are available on our user-friendly online knowledgebase, iModulonDB ^15^. Previously, we used 364 expression profiles of *P. aeruginosa* to generate and characterize 104 iModulons, which elucidated the genomic boundaries of biosynthetic gene clusters (BGCs), identified a novel bacteriocin producer, named ribosomally synthesized and post-translationally modified peptides (RiPP), that may play a role in virulence, and uncovered stimulons (clusters of iModulons with correlated activities) specific to iron and sulfur ^16^. Here, we expanded upon this TRN foundation by using new profiles generated from antibiotic treatment and biofilm growth conditions, since these are highly relevant for *P. aeruginosa* during infections of human hosts.

In this study, we used 411 high-quality expression profiles (281 publicly available in the SRA database and 130 generated for this study) to reconstruct the TRN of *P. aeruginosa* by implementing independent component analysis (ICA) using our established pipeline ^17^. We identified and characterized 116 iModulons comprising a quantitative TRN for *P. aeruginosa* based on all publicly available, high-quality RNA-sequencing profiles. Our analysis revealed novel insights into pathogenicity and antibiotic responses in this organism. We describe several two-component systems (TCSs) that regulate antibiotic efflux pumps, the differential activity of beta-lactam antibiotics in bacteriological and physiological media, and the influence of beta-lactam antibiotics on the Cell Division iModulon. To facilitate the broad utilization of this informative TRN, all iModulons can be searched, browsed, and studied through iModulonDB.org.

## Results

### Overview of *Pseudomonas aeruginosa* iModulons

In our previous study, we used 364 transcriptional profiles (281 from literature + 83 lab-generated) of *P. aeruginosa*, which is non-informative about the antibiotics responses ^16^. The aeruPRECISE364 profiles included the data from conditions like media types with different nutrient sources, osmotic stress, and specific gene-knockouts to identify the TRN associated with biosynthetic gene clusters, secretion systems, and central metabolism^16^. As the antibiotic specific responses are lacking in the previous study, we generated and compiled the antibiotic conditions in the current study.

The present study expands our transcriptome compendium from 364 to 411 high-quality profiles, introducing 47 new conditions including different antibiotic treatments and biofilm growth (**Supplementary Table S1, Figure 1a**). All transcriptomic profiles were processed using our established pipeline,^17^ all samples passed quality control, and all the replicates among the 411 profiles show Pearson’s correlation coefficient (PCC) of 0.98. To remove batch effects, all samples were normalized to respective reference conditions^17^. We applied our ICA workflow on the expanded compendium to compute an updated TRN composed of 116 independently modulated sets of genes, called iModulons (**Supplementary Figure S1b**). The new decomposition explains 66.16% of the total variance in the compendium of 411 profiles. Each iModulon was annotated with one of 11 functional categories: antibiotic resistance, amino acid and nucleotide metabolism, biosynthetic gene clusters, carbon source utilization, metal homeostasis, prophages, quorum sensing, stress response, structural components, translational, and uncharacterized.

**Figure 1.**
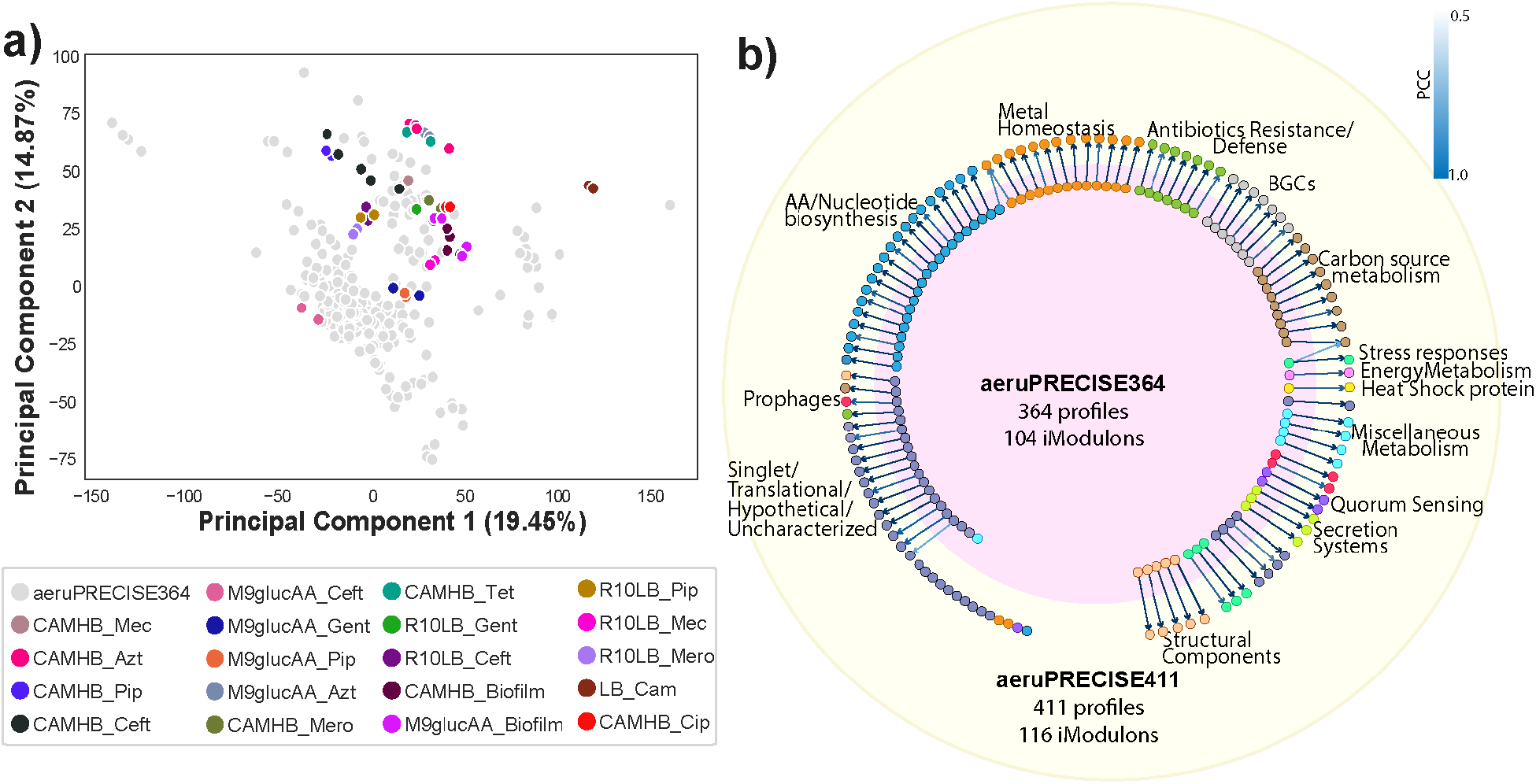
Overview of the iModulons identified in the aeruPRECISE411 study. a) Scatter plot showing the principal components of the 411 samples used in the study. The light gray colored dots represent the 364 samples used in the previous aeruPRECISE364 study ^16^, while the colored dots represent the new conditions, for antibiotic exposure and biofilm formation, new to this study. b) Comparison map showing the similarity and differences between the iModulons found in the aeruPRECISE364 ^16^ and aeruPRECISE411 (current) studies.

The transcriptional regulators of each iModulon were identified by comparing them with the known regulons. Regulon data was collected from RegPrecise^18^ and manually mined from literature (**Supplementary Table S1**). We used ‘regulon recall’ and ‘iModulon recall’ to characterize the association of the 116 iModulons with the known regulons. Regulon recall is the number of shared genes between an iModulon and its associated regulon divided by the total number of genes in the regulon, while the iModulon recall is the number of shared genes divided by the total number of genes in the iModulon. Based on the regulon recall and the iModulon recall and a threshold of 0.6, 116 iModulons are divided into 4 categories: well-matched, regulon subset, regulon discovery, and poorly matched (**Supplementary Figure S1b** and **Supplementary Notes 1**).

We compared the 116 iModulons identified in this study with our previous work^16^ that resulted in 104 iModulons (**Figure 1b** and **Supplementary Table S2**). By correlating the gene weights of the iModulons between the two datasets, we could map the previously characterized signals to the new ones. The large overlap between recovered signals indicates that the TRN structure is robust to new data, and the new iModulons show transcriptional response triggered by the new conditions used.

The new iModulons had functions relating to antibiotic resistance (AmpC), Metal Homeostasis (Fur and CysB-3), Amino Acid/Nucleotide biosynthesis (Lrp), Quorum sensing (Quorum sensing), Translation (Translational-3), Uncharacterized (Uncharacterized-14) and Singlets, which reflected the transcriptional signals likely to be activated by the newly added conditions. However, we found that for aeruPRECISE364, ArgR-2 (Amino Acid/Nucleotide biosynthesis) and Anr-1 (Metal homeostasis) merges into Anr-1 iModulon of the aeruPRECISE411 study, showing good correlation, with PCC of 0.78 and 0.61, respectively. Moreover, in aeruPRECISE364, RpoN (Carbon utilization) and RpoS-1 (Stress response) combine into the RpoN iModulon of aeruPRECISE411, with correlations of PCC 0.90 and 0.21, respectively ^16^(**Supplementary Figure S1d**).

### Role of two-component systems predicted in regulating efflux pumps

A two-component system (TCS) typically consists of a histidine kinase (HK) and a response regulator (RR)^19^. A TCS plays an important role in regulating various important functions, such as antibiotic resistance, virulence, and biofilm, among others ^20,21^. In the current study, we identified several TCSs that play a role in regulating efflux pumps. In total, *P. aeruginosa* has about 18 efflux pumps, which are responsible for increasing the resistance to numerous antibiotics ^22–24^.

We identified eight iModulons incorporating resistance-nodulation-cell division (RND) efflux pumps: *mexXY* (CzcR and MexZ), *czcABC* (CzcR), *mexCD-oprJ* (EsrC), *mexEF-oprN*(MexS-1), *mexGHI-opmD* (SoxR), *mexPQ-opmE* (CueR), *muxABC-opmB* (CpxR), and PA1874-PA1877 (PprB) (**Figure 2a, Table 1, and Supplementary Table S3**). The remaining 11 efflux pumps were not transcriptionally activated under the conditions in the dataset, and therefore were not identified by ICA. We identified seven RRs with strong correlations to the efflux-pump containing iModulons, indicating that they may be modulating the efflux pumps (**Figure 2a, red arrows**). Among them, we predicted potentially new RRs of the TCSs such as *mexT* (PA2492), PA2274, and PA3205 modulating the *mexEF-oprN, mexGHI-opmD,* and *muxABC-opmB* efflux pumps. Housseini *et al,* reported only five TCSs associated with RND efflux pumps were previously identified: RocS2-RocA2, ParR-ParS, AmgR-AmgS, CzcR-CzcS, and CopR-CopS ^25^. Therefore, our study uncovered new putative associations between TCSs and efflux pumps, with a relatively high correlation between expression of the identified RR and activity of the iModulon (**Figure 2c**). For example, our results suggest that the well studied CpxA-CpxR TCS regulates muxABC-opmB in response to beta lactams as part of the cell envelope stress response. We also used the activity levels of the iModulons under various conditions to identify metabolites which may activate the efflux pump containing iModulons, some of which were not known in the literature: DADS (EsrC), metals and organic compound (PprB), and n-acetyl-glucosamine (GlcNAc) (MexS-1) (**Figure 2b and Table 1**). Additional details on each efflux pump can be found in the **Supplementary Notes 2**.

**Figure 2.**
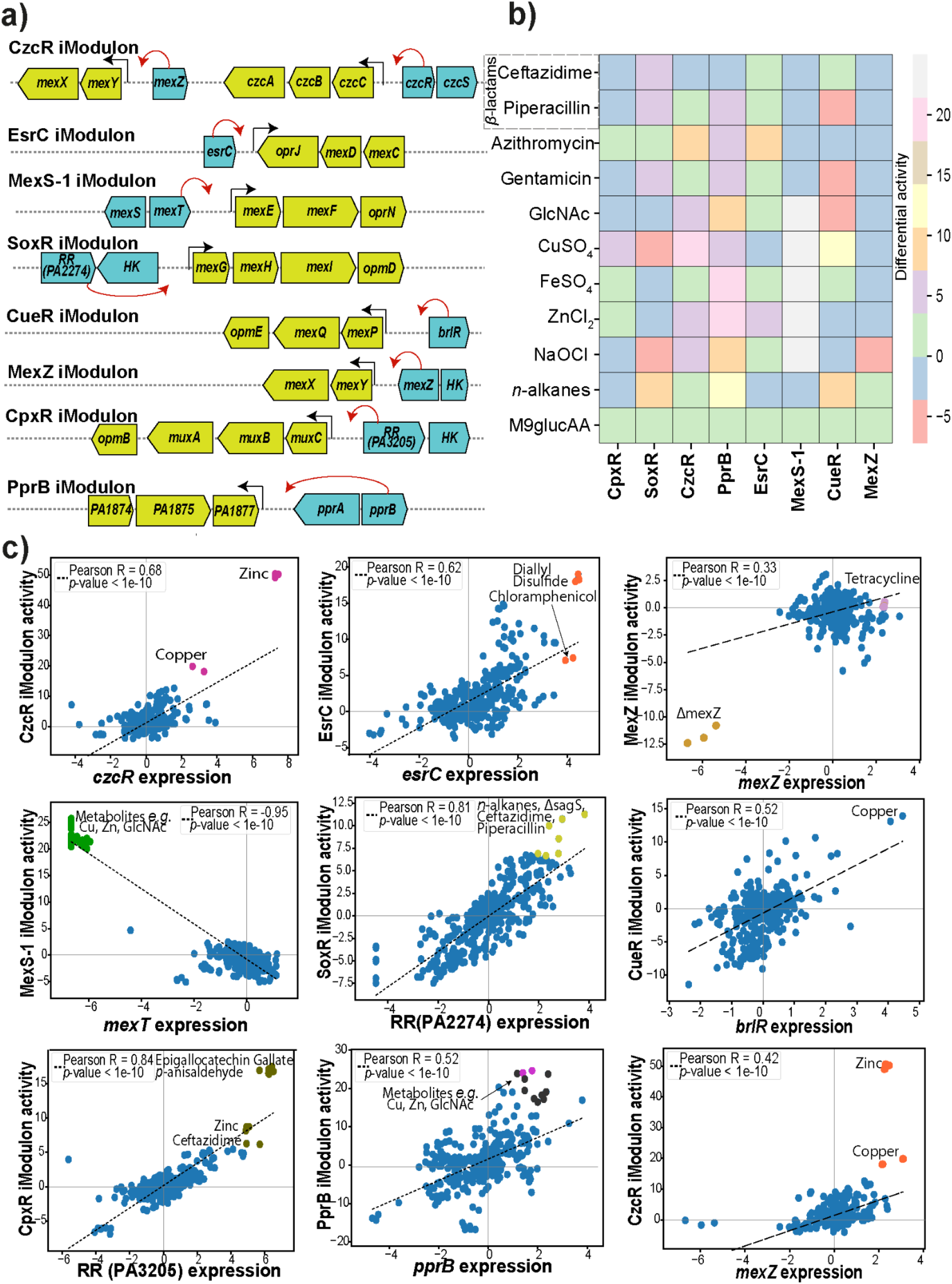
Role of Two-component systems (TCSs) in regulating efflux pumps in P. aeruginosa. a) Arrow diagram depicting the efflux pumps and their respective response regulator (RR, red arrow) of the TCS identified by our algorithm. b) Heatmap depicting the activity of iModulons containing efflux pumps under different treatments or growth conditions relative to the M9glucAA control. c) Scatter plot showing the correlation between the activity of efflux pump iModulons and the expression of their respective predicted RR genes. Colored dots indicate sampling conditions that activate the iModulons with regard to regulator genes.

**Table 1.** List of the iModulons related to the efflux pumps along with their regulator (Two-component system), efflux pumps, correlation of iModulon genes with the regulator, and activating conditions.

| iMod name | Predicted regulator (TCSs) | Efflux pumps/iMod | New regulator | Correlation value wrt each regulator | Activating condition |
| --- | --- | --- | --- | --- | --- |
| CzcR | Fur/CzcR | CzcCBA, MexXYZ | No | 0.68, 0.42 | Zinc, Copper |
| EsrC | EsrC | MexCD-OprJ | No | 0.62 | Diallyl disulfide, Chloramphenicol |
| MexS-1 | MexS/MexZ | MexEF-OprN | Yes | -0.95 | Macromolecules (Copper, Zinc, N-acetylglucosamine) |
| SoxR | SoxR | MexGHI-OpmD | Yes | 0.81 | *n*-alkanes, Ceftazidime, Piperacillin |
| CueR | BrIR | MexPQ-OpmE | No | 0.52 | Copper |
| MexZ | MexZ | MexXYZ | No | 0.33 | Tetracycline |
| CpxR | CpxR/ParB | MuxABC-Opm B | Yes | 0.84 | Epigallocatechin Gallate, *p*-anisaldehyde, Ceftazidime |
| PprB | PprB | PA1874-77 | No | 0.52 | Macromolecules (Copper, Zinc, N-acetylglucosamine) |

Furthermore, we compared the activities of the identified iModulons containing efflux pumps under different treatments or growth conditions (**Figure 2b**). As expected, there was high activity in the CzcR and CueR iModulons in response to copper (CuSO_4_) supplementation ^26^. Of interesting note is the high level of activation of the MexS-1 iModulon during growth on GlcNAc, during trace metal supplementation, and during NaOCl treatment (**Figure 2b**). A few studies suggest the increased concentration of GlcNAc, and metals like Cu, Zn, Fe, etc. are indicators of increased pathogenicity in the host in cystic fibrosis (CF) patients ^27,28^. Further, the *mexEF-oprN* efflux pump is known to increase the virulence in various conditions including CF ^29,30^. Our study suggests that the *mexEF-oprN* efflux pump is specific for GlcNAc, Cu, Fe, and Zn. Thus, we hypothesize that *mexEF-oprN* might play a role in increasing virulence in CF conditions.

Therefore, ICA analysis of the *P. aeruginosa* transcriptome was helpful in the identification of three TCSs (CpxR, SoxR, and MexS-1) which potentially regulate expression of efflux pumps (*muxABC-opmB, mexGHI-opmD, and mexEF-OprN, respectively*). We also identified substrates which specifically activate each efflux pump (as detailed in the form of a heatmap **Figure 2b, Table 1**). Further studies guided by these suggested relationships could elucidate the mechanisms underlying cellular transport and ultimately be useful for the design of antimicrobial drugs.

### Differential activity of beta-lactam antibiotics between bacteriological and physiological media

Beta-lactam antibiotics are often used clinically against *P. aeruginosa* infections, typically in conjunction with beta-lactamase inhibitors, to combat mechanisms of resistance in the pathogen. Recent studies have demonstrated that bacterial sensitivity to antibiotics can be altered depending on the growth medium used ^31,32^. We determined the minimal inhibitory concentration (MIC) of a variety of antibiotics against an efflux-pump knockout strain of *P. aeruginosa* in different media types, including the classic susceptibility testing media, CAMHB, and the eukaryotic cell culture media, RPMI 1640 supplemented with 10% Luria Broth (R10LB). We found that three of the four beta-lactam antibiotics tested had higher MICs in R10LB compared to CAMHB, whereas two of the three protein synthesis inhibitors had lower MICs in R10LB relative to CAMHB (**Table 2**).

**Table 2.** Minimum Inhibitory Concentration (MIC) values of antibiotics against P. aeruginosa K2733 (PAO1 ΔmexB) in different media types.

| Antibiotics | MOA | M9glucAA | CAMHB | R10LB | CAMHB/R10LB |
| --- | --- | --- | --- | --- | --- |
| Ceftazidime | Cell wall | 1 | 28.8 | 1 | 28.80 |
| Mecillinam | Cell wall | 4 | 14.4 | 64 | 0.225 |
| Meropenem | Cell wall | 0.06 | 0.06 | 0.5 | 0.120 |
| Piperacillin | Cell wall | 0.5 | 0.25 | 64 | 0.004 |
| Azithromycin | Protein | 1.6 | 8 | 0.4 | 20.00 |
| Gentamicin | Protein | 0.25 | 1.8 | 0.25 | 7.20 |
| Tetracycline | Protein | 0.5 | 0.9 | 28.8 | 0.031 |
| Ciprofloxacin | DNA | - | 0.014 | - | - |

Because of the observed differences in MIC, we surveyed the identified iModulons to determine if there were media-specific differences in the transcriptional response to the four beta-lactam antibiotics. We found that activities for certain iModulons containing antibiotic resistance-related genes, such as efflux pumps, had differential activity that could potentially be linked to media type (**Figure 3a,b**). The iModulon SoxR, for example, contains the genes *mexGHI* and *ompD* (**Figure 3c**) and has upregulated activity during treatment with beta-lactam antibiotics in R10LB and downregulated activity in CAMHB (**Figure 3a**), which correlates with the observed differences in MIC. The difference in the MIC value of antibiotics in both media types were also found in our previous studies on *Staphylococcus aureus* ^33–35^. Similarly, there was increased activity of the iModulon CzcR, involved in divalent cation homeostasis ^36^, in R10LB compared to CAMHB (**Figure 3a**). CzcR has been shown to mediate antibiotic susceptibility by regulating the expression of OprD, which is a route of entry for carbapenems like meropenem ^36^. Interestingly, there seemed to be no difference in activities for iModulons CpxR, CueR, and MexZ, while activities for the iModulon AmpC were upregulated compared to untreated controls but highly variable (**Figure 3a**).

**Figure 3.**
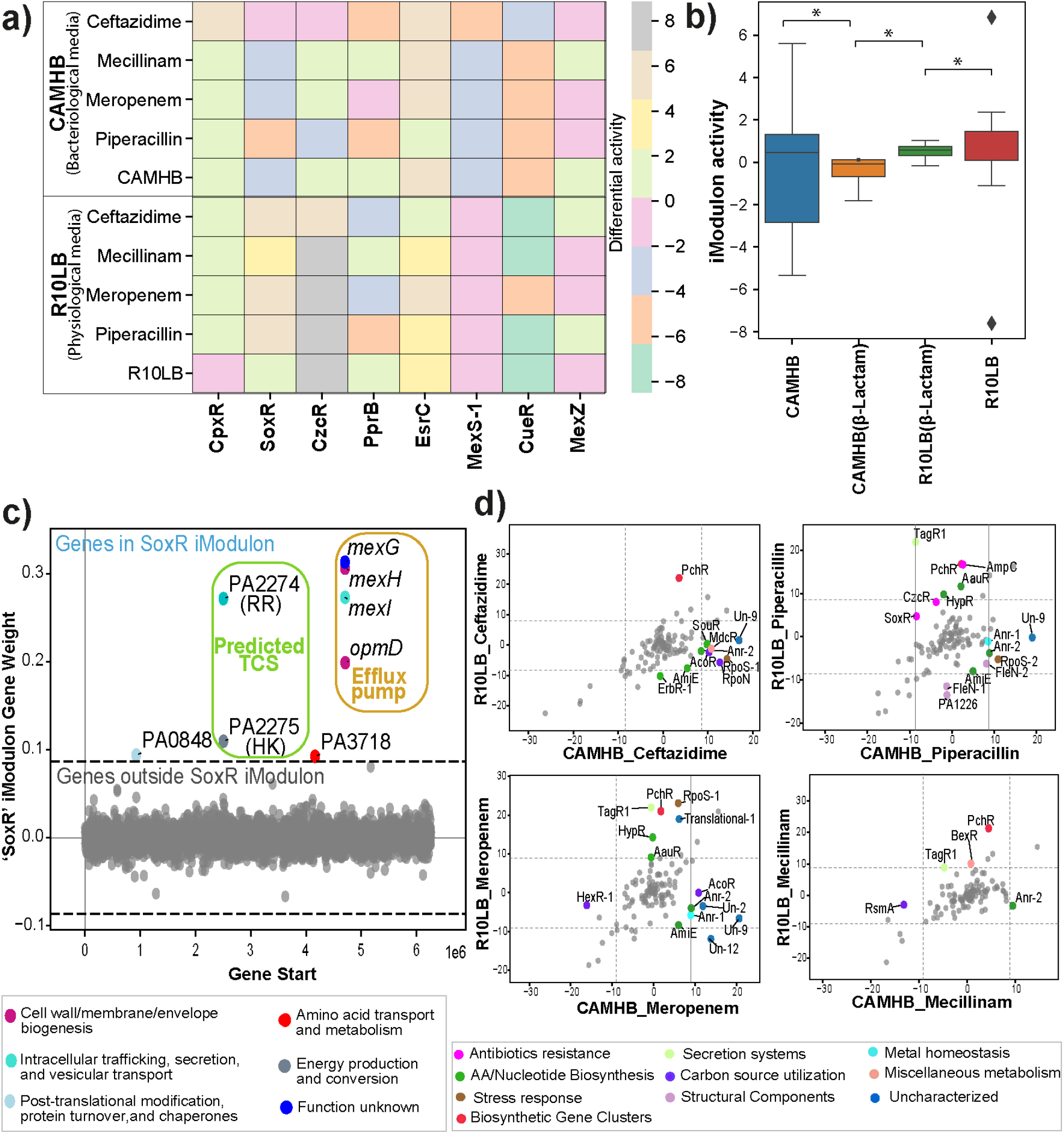
Differential activity of iModulons between the bacteriological and physiological media. a) Heatmap with the differential activity of the beta-lactam antibiotics (Ceftazidime, Mecillinam, Piperacillin, and Piperacillin) with respect to efflux pump-containing iModulons like CpxR, SoxR, CzcR, PprB, EsrC, MexS-1, CueR, MexZ, and AmpC. b) Boxplot showing the difference between the CAMHB and R10LB media for beta-lactam antibiotics in 8 efflux pumps (CpxR, SoxR, CzcR, PprB, EsrC, MexS-1, CueR, and MexZ). We find a significant difference in the activity of beta-lactam antibiotics among both media. c) Scatterplot showing the gene weights within the SoxR iModulon. The color of the dot corresponds to COG categories. d) Differential iModulon activity (DIMA) plot of the beta-lactam antibiotic treatments in CAMHB and R10LB. The color of the dots represents the functional category of iModulons.

When we compared all iModulon activities of individual beta-lactam treatments between CAMHB and R10LB, we found some media-specific differences. The iModulon TagR1 is upregulated in three of the four beta-lactam treatments in R10LB compared to CAMHB (**Figure 3d**). The TagR1 iModulon contains genes related to the type VI secretion system, specifically the PpkA protein kinase, which has been shown to mediate bacterial adaptability as well as oxidative stress tolerance and pyocyanin synthesis ^37,38^. This suggests that TagR1-mediated oxidative stress tolerance may help *P. aeruginosa* tolerate these antibiotics in the physiological milieu but not in CAMHB, where antibiotic susceptibility is typically tested. Additionally, the iModulon PchR, involved in pyochelin synthesis and iron acquisition ^39^, has high activity in R10LB conditions compared to the CAMHB samples (**Figure 3d**). In comparison, the iModulons Un-9, which encompasses genes in energy metabolism, and Anr-2, which contains genes related to iron acquisition, were both highly activated in the CAMHB conditions relative to the R10LB samples (**Figure 3d**). In combination, this data demonstrates that although the antibiotics have the same molecular targets within *P. aeruginosa*, the transcriptional response of the bacteria to the treatment can vary depending on media type.

### Differential activation of the Cell Division iModulon by beta-lactam antibiotics

Beta-lactam antibiotics have different affinities for various penicillin-binding proteins and enzymes responsible for coordinating cell wall synthesis and bacterial cell division ^40–42^. It has been shown that treatment of *Eshcerichia coli* with different antibiotics, including cell-wall active compounds, leads to morphologically distinct cytology based on molecular mechanisms of action ^43^. Mecillinam binds specifically to PBP-2, which is essential for the maintenance of rod-morphology in *P. aeruginosa, E. coli*, and other Gram-negative bacteria ^44^. Meropenem targets PBP-2 and PBP-4 ^41,45^, a non-essential DD-endopeptidase ^46,47^. Piperacillin and ceftazidime have specificity for PBP-3 ^41^, also known as FtsI, which is a member of the divisome and is important for the septation of daughter cells ^48,49^. *P. aeruginosa* treated with these antibiotics have morphological differences based on the PBP of the antibiotic targets (**Figure 4c**). Mecillinam-treated cells are smaller and more oblong-shaped, and penicillin or ceftazidime treatment leads to cell filamentation (**Figure 4c**). We were interested to see if the introduction of stress on the same biosynthetic pathway, i.e., cell wall, by different molecular means, in this case, beta-lactam antibiotics, would have the same or different transcriptional response in *P. aeruginosa*.

**Figure 4.**
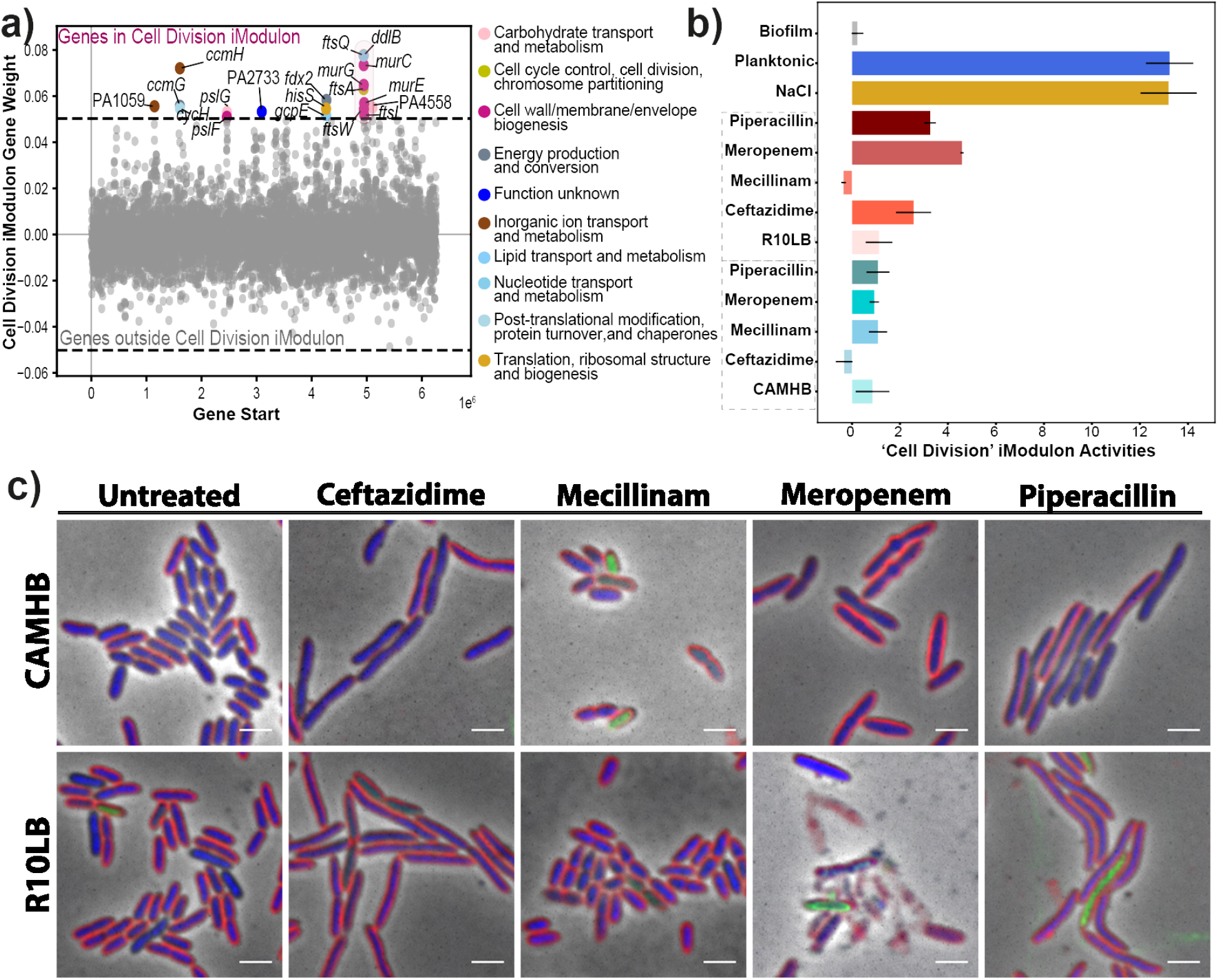
The Cell Division iModulon. a) Scatter Plot showing the gene weights within the Cell Division iModulon. The color of the dot corresponds to the COG categories. b) Bar plot depicting the Cell Division iModulon activities during treatment with beta-lactam antibiotics (Ceftazidime, Mecillinam, Piperacillin, and Piperacillin) in either CAMHB (bacteriological) or R10LB (physiological) media, along with the biofilm and planktonic (bioreactor) mode of growth. c) Microscopy images of the P. aeruginosa cells treated with beta-lactam antibiotics in either CAMHB or R10LB media. Cellular membranes are labeled with FM464 (red), DNA is labeled with DAPI (blue), and cellular permeability is indicated by the presence of the membrane-impermeable DNA dye SYTOX Green (green). The scale bar is 2 μm.

We identified a single Cell Division iModulon (**Figure 4a**). The Cell Division iModulon contained the genes *murC, murE*, and *murG*, which are important for the synthesis of peptidoglycan ^50–52^, as well as divisome components FtsA and FtsQ ^53,54^ (**Figure 4a**). This iModulon also contains PBP3, also known as FtsI (**Figure 4a**), which is the specific target for penicillin and ceftazidime ^41^. The Cell Division iModulon activity increased in R10LB compared to untreated but not when grown in CAMHB, during beta-lactam treatment, with the exception of mecillinam (**Figure 4b**). The untreated samples have similar levels of activity for this Cell Division iModulon (**Figure 4b**). Mecillinam treatment had little effect on the activity in this iModulon and also had a smaller effect on cell morphology. The higher levels of the Cell Division iModulon activity in the treated samples grown in R10LB may be a major contributor to the observed higher MICs in this media compared to CAMHB. The cells grown in R10LB produce more divisome related transcripts that likely allow for greater tolerance of cell wall-active antibiotic induced stress. The difference in MIC might also be due to the activation of efflux pumps, as noted earlier. We also discovered that there was high Cell Division iModulon activity in samples taken from planktonic growth (PRJNA643216) as well as in those grown in high salt conditions (**Figure 4b**), but low activity in samples taken from *P. aeruginosa* grown in a biofilm (PRJNA643216). This finding is confirmatory, since we would expect actively dividing cells, such as those growing planktonically in a bioreactor ^55^, to have high Cell Division iModulon activity compared to slower growing cells, such as those in a biofilm. Identifying the regulatory mechanisms underlying this iModulon’s activation should be the topic of further study, as additional drugs which downregulate this response could be administered in conjunction with beta-lactams to improve their effectiveness.

### The PprB iModulon activates under stress conditions and is a signal for the transition from planktonic to biofilm lifestyle

The PprB iModulon consists of the PA1874-PA1877 efflux pump, tight adherence (*tad*)machinery, and the *cup*E1-*cup*E5 (*flp* fimbriae) chaperone-usher pathway (**Figure 5a,b**). We found that the PprB iModulon is highly activated during biofilm growth in M9 media supplemented with glucose and amino acids (M9glucAA) and cation-adjusted Mueller-Hinton broth (CAMHB) conditions as well as during growth with iron (Fe), zinc (Zn), and *n*-alkane supplementation (**Figure 5c**). It has been previously established that PprB is involved in biofilm formation ^56,57^, including the regulation of *tad* machinery expression involved in the production of the Type IVb pili ^58,59^. Additionally, PprB was demonstrated to control fimbriae assembly *via* CupE, which is specifically expressed during biofilm growth ^60^. Although the molecules excreted by the efflux pump are unknown, the *tad* system and *cup*E1-E5 system both support biofilm formation. The activation of this set of genes under high-Fe conditions supports recent evidence that high-Fe conditions lead to a limited motility phenotype that promotes biofilm formation ^61^.

**Figure 5.**
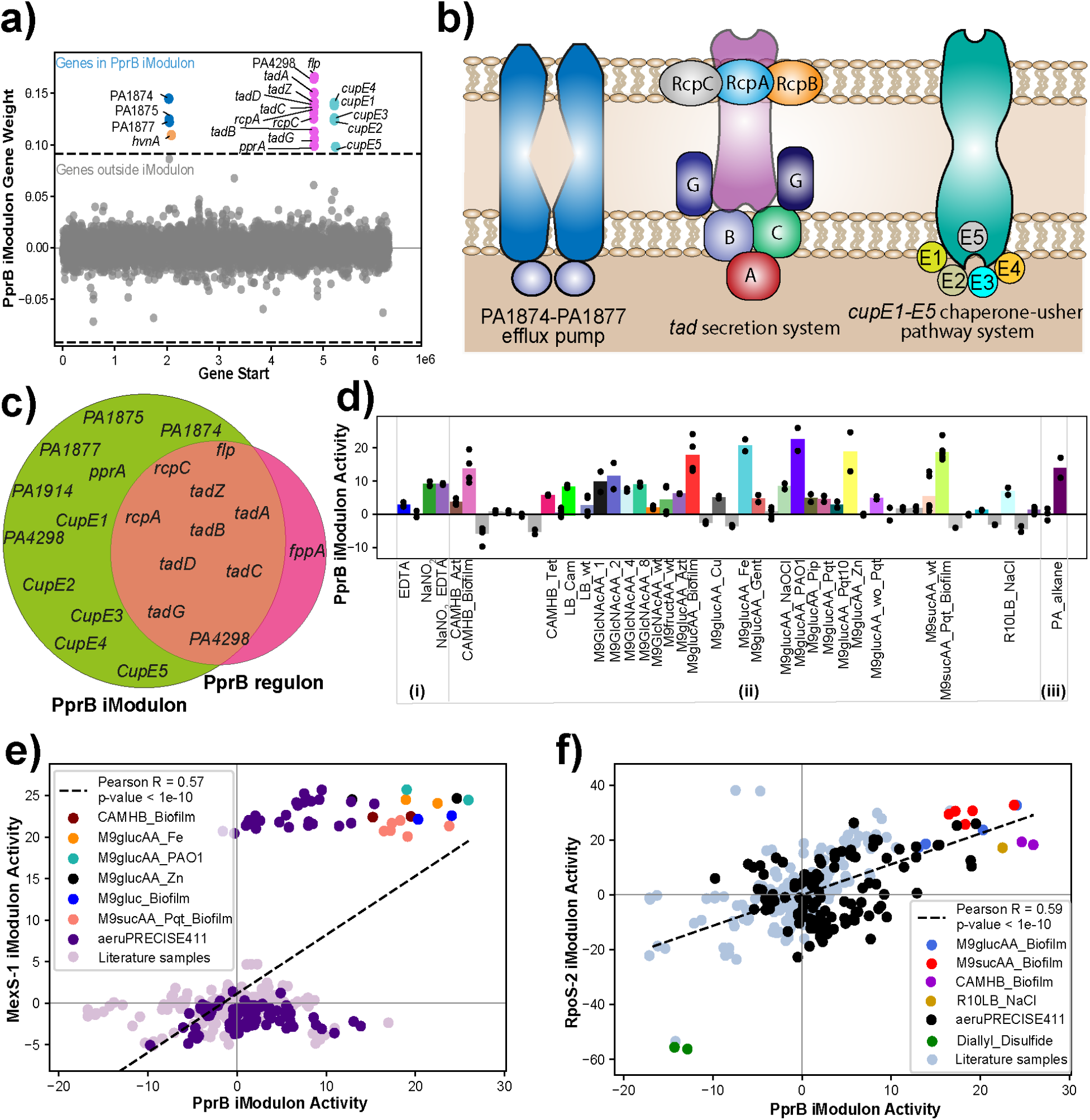
The iModulon PprB may be a signal for the transition to biofilm growth. a) Scatter Plot showing the gene weights of the PprB iModulon; the color depicts the COG categories. The PprB iModulon shows three components; PA1874-PA1877 efflux pump, tad machinery, and the cupE1-E5 chaperone-usher pathway system. b) Schematic diagram of the PprB iModulon depicting its three components. c) Venn diagram showing the genes in the PprB iModulon and PprB regulon. d) Activity plot of the conditions expressed in the PprB iModulon in the AB569_EDTA_NaNO2(i), aeruPRECISE411(ii), and n-alkanes_treatment (iii). e) Scatter plot showing the correlation (PCC 0.57) between the MexS-1 iModulon and the PprB iModulon. f) Scatter plot showing the correlation (PCC 0.59) between the RpoS-2 iModulon (stress-responsive) and the PprB iModulon.

The genes PA1874 through PA1877 were included in the PprB iModulon. This operon was previously characterized as a likely type I secretion system (T1SS), important for resistance to aminoglycosides and fluoroquinolones, specifically during growth in biofilms ^57,62^. Further, these studies observed the increased susceptibility to the aminoglycoside tobramycin during constitutive expression of *pprB* in biofilm growth, which was thought to be through *pprB* modulation of membrane permeability ^63^. We found that the PprB iModulon is activated during planktonic growth with treatment with antibiotics that target protein synthesis, such as gentamicin, azithromycin, and tetracycline, as well as NaOCl and paraquat (pqt), which are both redox active compounds (**Figure 2c, 5d**). This suggests that the PprB iModulon is activated under stress conditions and may be a contributing singal for *P. aeruginosa* to switch between planktonic and biofilm modes of growth.

We found significant correlation between the activity levels of the PprB iModulon and the MexS-1 and RpoS-1 iModulons. We found that the MexS-1 iModulon had a strong bimodal activity distribution, and all samples which highly activated the MexS-1 iModulon also activated the PprB iModulon. These samples included the presence of supplemental Fe and Zn, during growth on GlcNAc, as well as during the biofilm mode of growth (**Figure 5e, 5f**). We also found a linear correlation between the activities of the PprB and the RpoS-2 iModulons, the latter of which contains the central stress response sigma factor RpoS. Both iModulons activated during growth in biofilm or osmotic stress (e.g., NaCl), and were highly inactivated during treatment with diallyl disulfide (DADS) (**Figure 5f**) Treatment of *P. aeruginosa* with DADS was shown to down-regulate many genes involved in pathogenicity and virulence, including TCSs, such as PprB ^64^. These correlations indicate either co-stimulation or hierarchical regulation; in the case of RpoS-2, it suggests that stress correlates with biofilm formation, which is consistent with expectations about the function of biofilm in protecting *P. aeruginosa* communities. Stewart *et al* also shows that stress can contribute to antibiotic resistance in biofilms of *P. aeruginosa*.

## Discussion

*P. aeruginosa* is an important member of the ESKAPEE group of pathogenic bacteria ^65^. It is an opportunistic pathogen and shows high resistance towards diverse classes of antibiotics ^66^. In the current study, we used ICA on 411 expression profiles of *P. aeruginosa* to generate 116 iModulons. Our analysis reveals various important aspects of *P. aeruginosa* transcriptional regulation, such as prediction of TCSs regulating the efflux pumps, specific macromolecules and antibiotics which activate transcription of TCS-dependent efflux pumps, differential behavior of beta-lactam antibiotics in bacteriological and physiological media, impact of beta-lactam antibiotics on the PBPs and cell division, and the role of the PprB iModulon in the transition to biofilm growth under stress conditions.

The expression of efflux pumps is one of the classic mechanisms of antibiotic resistance ^67^. Apart from antibiotics, efflux pumps transport various compounds like metals and organic compounds out of the cell ^68^. In this study, we predicted the RRs of TCSs which may regulate efflux pumps based on correlated transcription (**Figure 2 and Table 1**). Further, we used the activating conditions of the efflux pumps to predict their substrates, which identified new putative relationships.

We found that antibiotics sharing the same molecular target can activate iModulons for efflux pumps or cell division to different levels, and their activities are also dependent on the media type in which the bacteria are (**Figure 3, 4**). Therefore, this work provides a molecular view into transcriptional regulation underpinning the media-specific susceptibility differences observed via MIC between *P. aeruginosa* treated in the bacteriological media CAMHB and the LB-supplemented eukaryotic cell culture media R10LB (**Table 2**).

We also identified a single cell division iModulon of *P. aeruginosa* that had differential activity dependent on media condition, e.g., CAMHB vs R10LB, and mode of growth, e.g., biofilm vs planktonic. Confirming experimental observations of higher MICs during growth in R10LB, we found an increased activity of the cell division iModulon during beta-lactam antibiotic treatment in R10LB relative to untreated and to CAMHB conditions. This highlights the ability of iModulons to identify potentially clinically-relevant transcriptional responses from *in vitro* experimental data sets.

The current study is focused on elucidating the antibiotic-specific behavior of *P. aeruginosa* through the application of machine learning to a compendium of transcriptomic profiles. The code used to explore the iModulons is available at GitHub (https://github.com/akanksha-r/modulome_paeru2.0). To make our data easily accessible to experimental biologists we provide our data in the form of a user-friendly web interface (https://imodulondb.org/dataset.html?organism=P_aeruginosa&dataset=precise411). We found that the iModulons are an important tool for the reconstruction of the TRN of *P. aeruginosa* to address various biological queries, including predicting the regulators and substrates of efflux pumps, examining the differential behavior of beta-lactam antibiotics in different media, exploring the role of biofilm-specific antibiotic resistance systems in the virulence of *P. aeruginosa*, and elucidating the molecular mechanism of beta-lactam antibiotics on PBPs and cell division. This study demonstrates that iModulon-based TRNs can provide new insights when new transcriptomic data is added; this work can therefore serve as the basis for further studies. With increasing scale, we hope that this approach will culminate in a comprehensive, quantitative, irreducible TRN for this important pathogen.

## Methods

### RNA extraction and library preparation

*P. aeruginosa* (PAO1 and *ΔmexB*) strains were used in this study. We extracted RNA samples for 45 unique conditions including different media types (M9, CAMHB, LB, RPMI+10%LB), oxidative stress (treatment with paraquat), iron starvation (treatment with DPD), osmotics stress (high NaCl), low pH, various carbon sources (succinate, glycerol, pyruvate, fructose, sucrose, N-acetyl glucosamine), micronutrients (copper, iron, zinc, sodium hypochlorite), antibiotics (gentamicin, azithromycin, piperacillin, ceftazidime, mecillinam, meropenem, tetracycline), and biofilms. All conditions were collected in biological duplicates and untreated controls were also collected for each set to rule out the possibility of the batch effect (**Supplementary Table S1**).

We followed the protocol outlined by Rajput *et al* to perform RNA extraction and library preparation ^16^. In brief, strains were grown overnight at 37°C, with rolling, in appropriate media types for the testing condition of choice. Overnight cultures were then diluted to a starting OD_600_ of ~0.01 and grown at 37°C, with stirring. Once cultures reached the desired OD_600_ of 0.4, 2 mL cultures were immediately added to centrifuge tubes containing 4 mL RNAprotect Bacteria Reagent (Qiagen), vortexed for 5 seconds and incubated at room temperature for 5 min. Samples were then centrifuged for 10 minutes at 5000xg and the supernatant was removed prior to storage at −80°C until further processing. In conditions involving antibiotic treatment, when the bacterial culture had reached an OD_600_ of ~0.2, antibiotics were added at 2X or 5X their MIC in the appropriate media type and allowed to incubate at 37°C, with stirring, for an additional hour prior to sample collection.

Total RNA was isolated and purified using a Zymo Research Quick-RNA Fungal/Bacterial Microprep Kit from frozen cell pellets previously harvested using Qiagen RNAprotect Bacteria Reagent according to the manufacturers’ protocols. Ribosomal RNA was removed from 1 ug Total RNA with the use of a thermostable RNase H (Hybridase) and short DNA oligos complementary to the ribosomal RNA, performed at 65 degrees C to prevent non-specific degradation of mRNA. The resulting rRNA-subtracted RNA was made into libraries with a KAPA RNA HyperPrep kit incorporating short Y-adapters and barcoded PCR primers. The libraries were quantified with a fluorescent assay (dsDNA AccuGreen quantitation kit, Biotium) and checked for proper size distribution and average size with a TapeStation (D1000 Tape, Agilent). Library pools were then assembled and a 1X SPRI bead cleanup performed to remove traces of carryover PCR primers. The final library pool was quantified and run on an Illumina instrument (NextSeq, Novaseq).

### Fluorescence Microscopy

After collection of RNAseq samples, fluorescence microscopy was performed as previously described ^43^. 4 μL cells were added to 4 μL dye mix containing 100 μg/mL FM4-64 (red), 100 μg/mL DAPI (blue), and 20 μM SYTOX Green (green) in 1x T-base. Samples were transferred to a single-well glass slide containing an agarose pad (1.2% agarose in 20% LB) and imaged on an Applied Precision DV Elite epifluorescence microscope. The exposure times for each wavelength were kept constant for all images.

### Data preprocessing

Apart from the in-house generated data, we also downloaded and processed all RNA sequencing data available from NCBI SRA for *P. aeruginosa* PAO1. In the current study, we used 411 expression profiles, in which 281 is collected from the NCBI-SRA and 130 is generated in the lab. Out of 130 lab generated expression profiles, 83 includes the conditions like osmotic stress, carbon sources (glucose, succinate, pyruvate, glycerol, N-acteyglucosamine), metals (Zn, Cu, Fe), media types (CAMHB, M9, and R10LB) and salt stress, which are used in our previous study. However, in the current study, we added some conditions like antibiotics and biofilms to reconstruct the TRNs. We include antibiotics like gentamicin, azithromycin, piperacillin, ceftazidime, mecillinam, meropenem, and tetracycline in media like CAMHB, R10LB, and M9.

Data processing and quality control for the public datasets is detailed in Sastry *et al* 2021 ^17^. Data processing and quality control scripts are available at https://github.com/avsastry/modulome-workflow. Briefly, raw FASTQ files were downloaded from NCBI using fasterq-dump (https://github.com/ncbi/sra-tools/wiki/HowTo:-fasterq-dump). Next, read trimming was performed using Trim Galore (https://www.bioinformatics.babraham.ac.uk/projects/trim_galore/) with the default options, followed by FastQC (http://www.bioinformatics.babraham.ac.uk/projects/fastqc/) on the trimmed reads. Next, reads were aligned to the *P. aeruginosa* genome (NC_002516.2) using Bowtie ^69^. The read direction was inferred using RSEQC ^70^ before generating read counts using featureCounts ^71^. Finally, all quality control metrics were compiled using MultiQC ^72^ and the final expression dataset is reported in units of log-transformed Transcripts per Million (log-TPM).

To ensure quality control, data that failed any of the following four FASTQC metrics were discarded: per base sequence quality, per sequence quality scores, per base n content, and adapter content. Samples that contained under 500,000 reads mapped to coding sequences were also discarded. Hierarchical clustering was used to identify samples that did not conform to a typical expression profile.

Manual metadata curation was performed on the data that passed the first four quality control steps. Information including the strain description, base media, carbon source, treatments, and temperature were pulled from the literature. Each project was assigned a short unique name, and each condition within a project was also assigned a unique name to identify biological and technical replicates. After curation, samples were discarded if (a) metadata was not available, (b) samples did not have replicates, or (c) the Pearson R correlation between replicates was below 0.95. Finally, the log-TPM data within each project was centered to a project-specific reference condition. After quality control, the final compendium contained 411 high-quality expression profiles: 130 generated for this study, plus 281 expression profiles extracted from public databases.

### Computing robust component with ICA

To compute the optimal independent components, an extension of ICA was performed on the RNA-seq dataset as described in McConn *et al* ^73^.

Briefly, the scikit-learn (v0.23.2) ^74^ implementation of FastICA ^75^ was executed 100 times with random seeds and a convergence tolerance of 10^-7^. The resulting independent components (ICs) were clustered using DBSCAN ^76^ to identify robust ICs, using an epsilon of 0.1 and minimum cluster seed size of 50. To account for identical components with opposite signs, the following distance metric was used for computing the distance matrix:

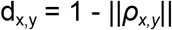

where *ρ_x,y_* is the Pearson correlation between components *x* and *y*. The final robust ICs were defined as the centroids of the cluster.

Since the number of dimensions selected in ICA can alter the results, we applied the above procedure to the dataset multiple times, ranging the number of dimensions from 10 to 360 (i.e., the approximate size of the dataset) with a step size of 10. To identify the optimal dimensionality, we compared the number of ICs with single genes to the number of ICs that were correlated (Pearson R > 0.7) with the ICs in the largest dimension (called “final components”). We selected the number of dimensions where the number of non-single gene ICs was equal to the number of final components in that dimension.

### Determination of the gene coefficient threshold

The gene coefficients are determined as described in Sastry *et al* ^17^. Each independent component contains the contributions of each gene to the statistically independent source of variation. Most of these values are near zero for a given component. In order to identify the most significant genes in each component, we iteratively removed genes with the largest absolute value and computed the D’Agostino K^2^ test statistic ^77^ for the resulting distribution. Once the test statistic dropped below a cutoff, we designated the removed genes as significant.

To identify this cutoff, we performed a sensitivity analysis on the concordance between significant genes in each component and known regulons. First, we isolated the 20 genes from each component with the highest absolute gene coefficients. We then compared each gene set against all known regulons using the two-sided Fisher’s exact test (FDR < 10^-5^). For each component with at least one significant enrichment, we selected the regulator with the lowest p-value.

Next, we varied the D’Agostino K^2^ test statistic from 50 through 2000 in increments of 50, and computed the F1-score (harmonic average between precision and recall) between each component and its linked regulator. The maximum value of the average F1-score across the components with linked regulators occurred at a test statistic of cutoff of 420 for the *P. aeruginosa* dataset.

For future datasets where a draft TRN is unavailable, an alternative method is proposed that is agnostic to regulator enrichments. The Sci-kit learn ^74^ implementation of K-means clustering, using three clusters, can be applied to the absolute values of the gene weights in each independent component. All genes in the top two clusters are deemed significant genes in the iModulon.

### iModulon annotation

The regulator enrichments are determined as described in Sastry *et al* ^17^. The gene annotation pipeline can be found at https://github.com/SBRG/pymodulon/blob/master/docs/tutorials/creating_the_gene_table.ipynb. Gene annotations were pulled from Pseudomonas genome db ^5^. Additionally, KEGG ^78^ and Cluster of Orthologous Groups (COG) information were obtained using EggNOG mapper ^79^. Uniprot IDs were obtained using the Uniprot ID mapper ^80^, and operon information was obtained from Biocyc ^81^. Gene ontology (GO) annotations were obtained from AmiGO2 ^82^. The known TRN was obtained from RegPrecise ^18^ and manually curated from literature.

### Differential activation analysis

The gene coefficients are determined as described in Rajput *et al* ^16^. We calculated the difference in the iModulons activities between two or more conditions by fitting the log-normal distribution to the difference. The statistical significance was checked by calculating the difference between the absolute values of the activities among the iModulons, and further confirmed by the *p*-values. However, the *p*-values were adjusted using the Benjamini-Hochberg correction. While calculating the differential activation among the iModulons, the difference with >5 was considered significant.

### Prediction of two-components system

Through our iModulon analysis, we identified the TCSs regulating the efflux pumps. It is done in 3 steps: 1) Identification of the TCSs by using our previously published method ^20^ and P2CS database ^83^. 2) Checking the relationship between the RR gene and the iModulon containing efflux pumps by calculating the PCC between them. 3) Validating the RR’s specificity/relationship by scanning its correlation with all predicted 116 iModulons. Furthermore, we also checked the specificity of the metabolites with respective efflux pumps.

### Generating an iModulonDB web page

The *P. aeruginosa aeru*PRECISE411 iModulonDB ^15^ web pages was generated using the imodulondb_export() function in the Pymodulon package ^17^. The generated page includes gene information from Pseudomonas Genome DB (https://pseudomonas.com)^5^.

## Data availability

All the in-house generated sequences were deposited in the NCBI-Sequence Read Archive database (PRJNA717794). The accession number of the deposited reads is provided in the **Supplementary Table S1**. While the X, M and A matrices are available at GitHub (https://github.com/akanksha-r/modulome_paeru2.0). Each gene and iModulon have interactive, searchable dashboards on iModulonDB.org, and data can also be downloaded from there.

## Code availability

The customized code for the ICA analysis as well as various files including the X, M, A matrices, TRN regulator file, gene annotated files, gene ontology and kegg pathway annotation files are available on GitHub (https://github.com/akanksha-r/modulome_paeru2.0).

## Author contributions

A.R., A.V.S., J.P. and B.O.P. designed research. A.R. performed research. H.T. collected the experimental data. R.S. performed RNA isolation and library construction. K.R. developed a web interface. A.R., H.T., A.V.S., R.S., K.R. and S.M.C. performed the analysis. A.R. and H.T. wrote the manuscript. All the authors have read and approved the manuscript.

## Acknowledgments

We thank Marc Abrams for reviewing the manuscript and providing constructive suggestions. This work was supported by NIH Grant U01 AI124316 and Novo Nordisk Foundation Grant NNF20CC0035580.

