## Supplementary Figure S1, Supplementary Figure S2, Supplementary Figure S3, Supplementary Figure S4 for "Advanced transcriptomic analysis reveals the role of efflux pumps and media composition in antibiotic responses of *Pseudomonas aeruginosa*"

### Supplementary Information

#### Supplementary Figures

***Supplementary Figure S1****. Overview of aeruPRECISE411 iModulons. a) Schematic diagram showing the overview of ICA to calculate independently modulated sets of genes named iModulons. The expression profiles (X) are decomposed into M and A matrices. The M matrix represents the independent components as a set of genes, while the A matrix shows the condition-specific activities. b) Schematic diagram depicting the iModulon recall and regulon recall calculation. c) Quadrant plot shows the four categories of iModulons i.e., Well-matched, Regulon Subset, Poorly-matched, and Regulon Discovery. The size of the circles represents the corresponding iModulon's explained variance. d) Cluster map depicting the passed samples in each cluster.*

***Supplementary Figure S2****. Role of Two-component system in regulating efflux pumps. a) Line plot depicting the correlation (Pearsons' correlation coefficient, PCC) between the expression of* *mexZ gene and 116 iModulons. b)  Line plot depicting the correlation (PCC) between the expression of brlR gene and 116 iModulons.*

***Supplementary Figure S3***. *Role of Two-component system in regulating efflux pumps. a) Line plot depicting the correlation (Pearsons' correlation coefficient, PCC) between the expression of czcR gene and 116 iModulons. b) Line plot depicting the correlation (PCC) between the expression of esrC gene and 116 iModulons. c)  Line plot depicting the correlation (PCC) between the expression of PA2274 gene and 116 iModulons.*

***Supplementary Figure S4***. *Role of Two-component system in regulating efflux pumps. a) Line plot depicting the correlation (Pearsons' correlation coefficient, PCC) between the expression of mexT gene and 116 iModulons. b) Line plot depicting the correlation (PCC) between the expression of PA3205 gene and 116 iModulons. c)  Line plot depicting the correlation (PCC) between the expression of pprB gene and 116 iModulons.*

#### Supplementary Notes

1) Categories of iModulons
2) Efflux Pumps

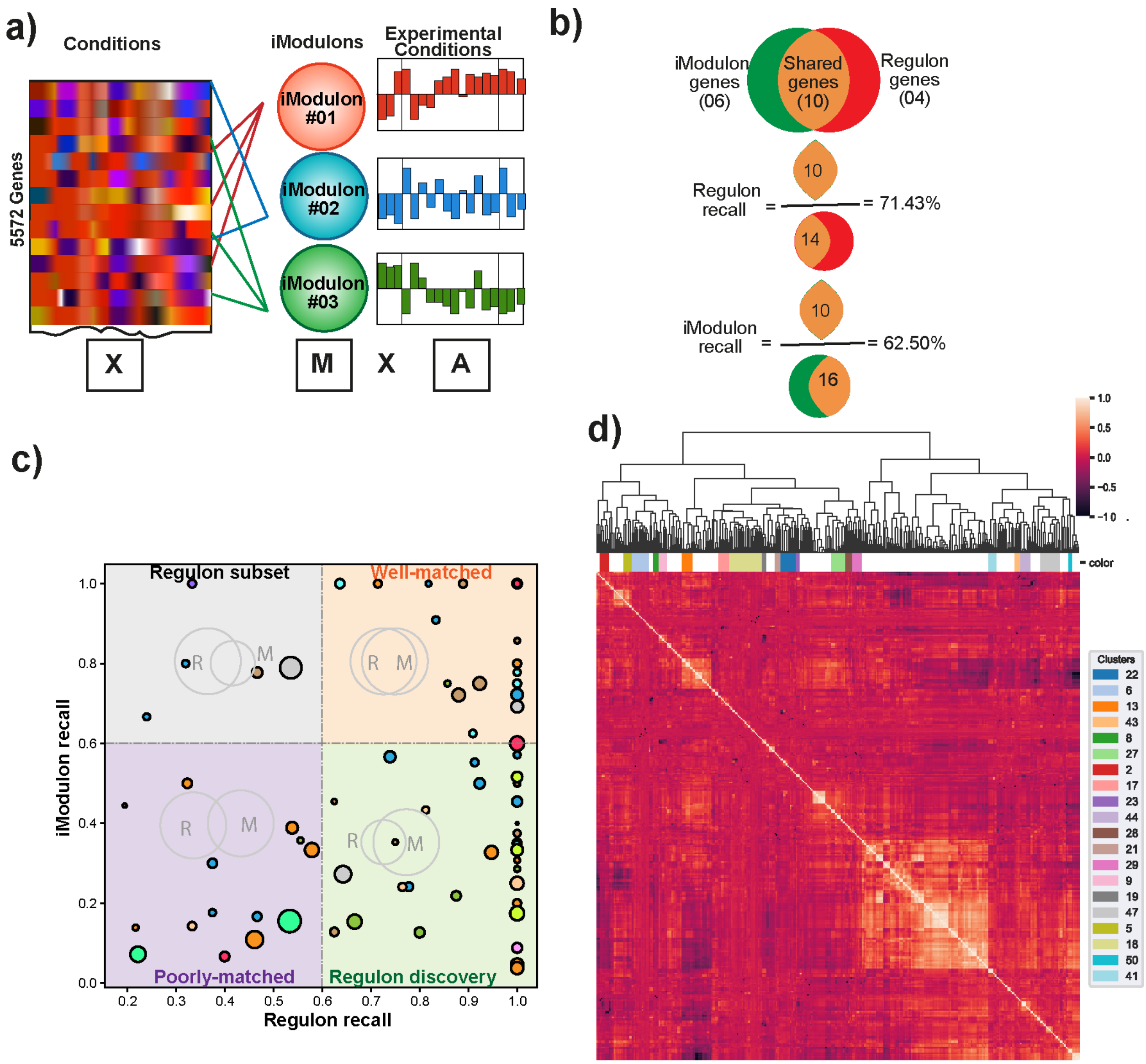


***Supplementary Figure S1***. *Overview of aeruPRECISE411 iModulons. a) Schematic diagram showing the overview of ICA to calculate independently modulated sets of genes named iModulons. The expression profiles (X) are decomposed into M and A matrices. The M matrix represents the independent components as a set of genes, while the A matrix shows the condition-specific activities. b) Schematic diagram depicting the iModulon recall and regulon recall calculation. c) Quadrant plot shows the four categories of iModulons i.e., Well-matched, Regulon Subset, Poorly-matched, and Regulon Discovery. The size of the circles represents the corresponding iModulon's explained variance. d) Cluster map depicting the passed samples in each cluster.*

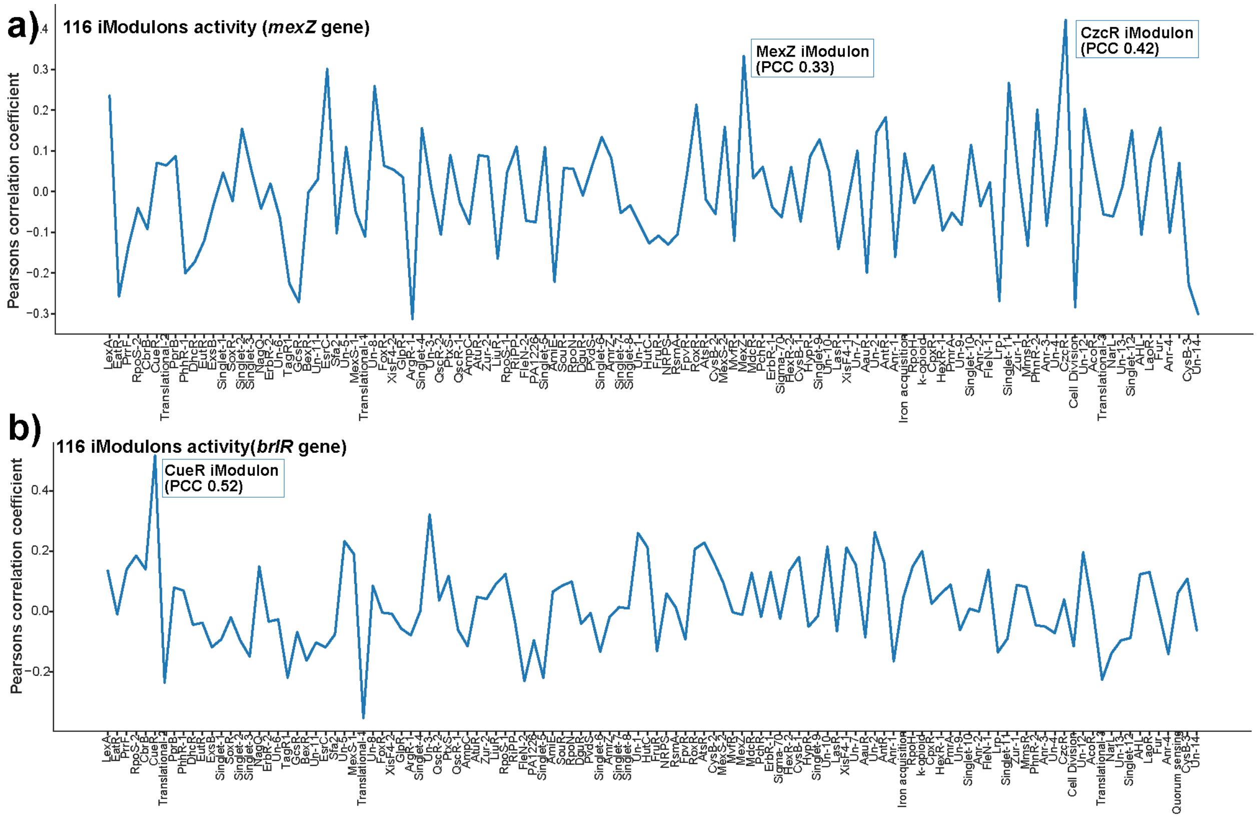


***Supplementary Figure S2***. *Role of Two-component system in regulating efflux pumps. a) Line plot depicting the correlation (Pearsons' correlation coefficient, PCC) between the expression of mexZ gene and 116 iModulons. b) Line plot depicting the correlation (PCC) between the expression of brlR gene and 116 iModulons.*

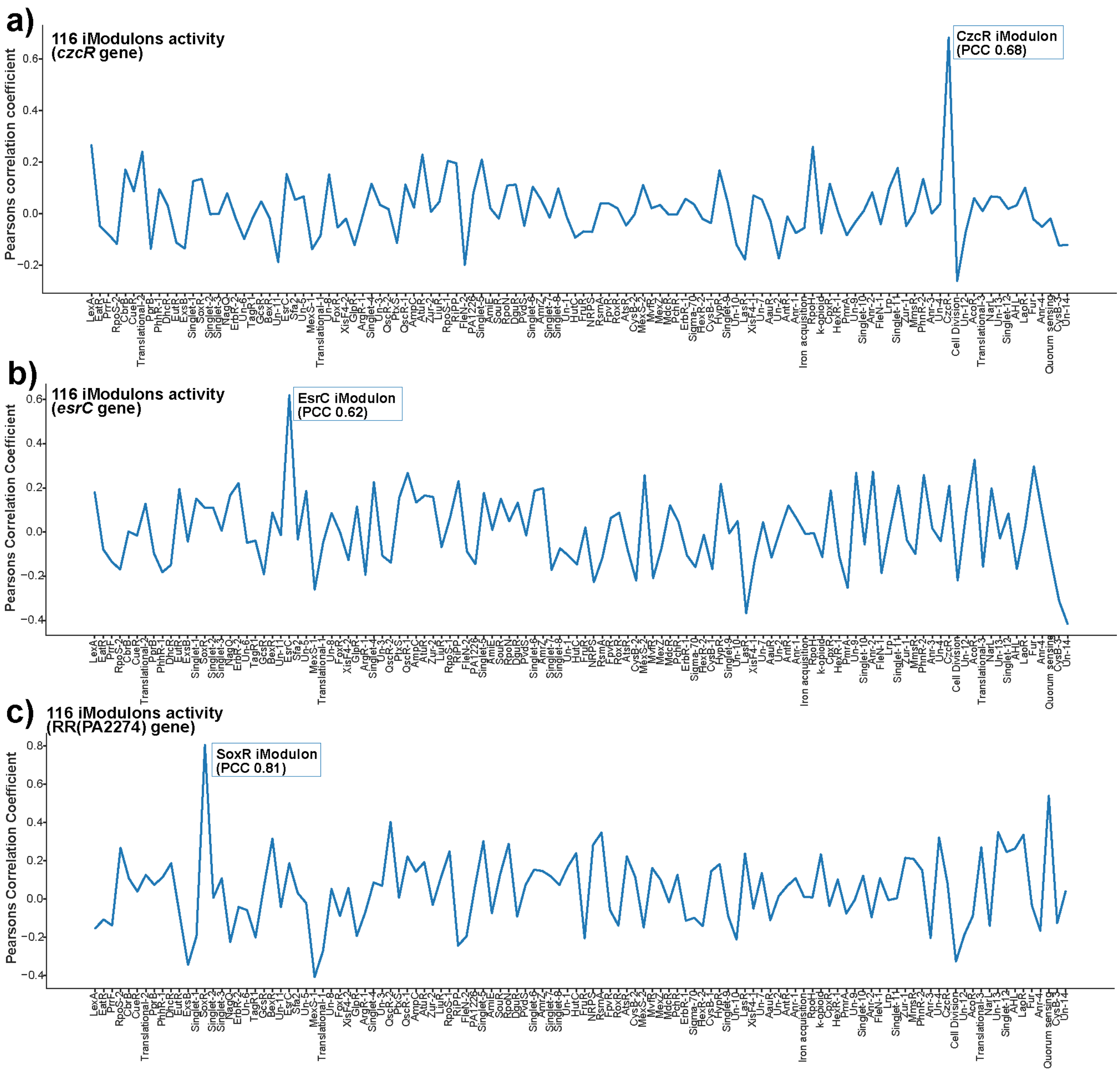


***Supplementary Figure S3***. *Role of Two-component system in regulating efflux pumps. a) Line plot depicting the correlation (Pearsons' correlation coefficient, PCC) between the expression of czcR gene and 116 iModulons. b) Line plot depicting the correlation (PCC) between the expression of esrC gene and 116 iModulons. c) Line plot depicting the correlation (PCC) between the expression of PA2274 gene and 116 iModulons.*

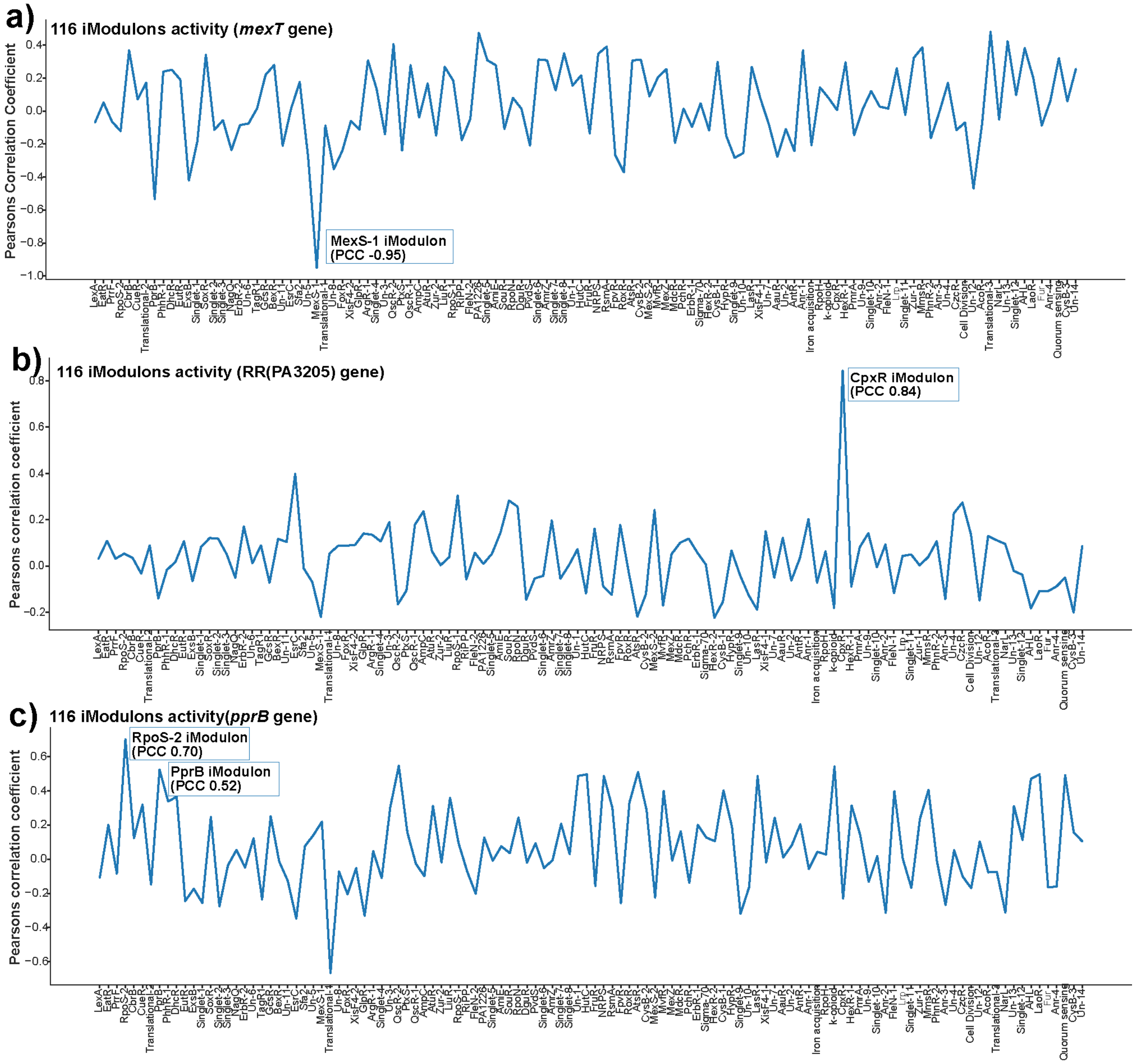


***Supplementary Figure S4****. Role of Two-component system in regulating efflux pumps. a) Line plot depicting the correlation (Pearsons' correlation coefficient, PCC) between the expression of mexT gene and 116 iModulons. b) Line plot depicting the correlation (PCC) between the expression of PA3205 gene and 116 iModulons. c) Line plot depicting the correlation (PCC) between the expression of pprB gene and 116 iModulons.*

#### Supplementary Notes

3) **Categories of iModulons**
4) **Efflux Pumps**

##### 1. Categories of iModulons

Based on the regulon recall and the iModulon recall and a threshold of 0.6, 116 iModulons are divided into 4 categories: well-matched, regulon subset, regulon discovery, and poorly matched (**Supplementary Figure S1b**). 1) The well-matched group (upper right quadrant) includes iModulons with regulon recall and iModulon recall greater than 0.60. The genes in these iModulons correspond to the existing regulons. 2) The regulon subset of iModulons in the upper left quadrant has high iModulon recall and low regulon recall, thus these iModulons represent only a part of a defined regulon. 3) The regulon-discovery iModulons (lower right quadrant) have high regulon recall and low iModulon recall. The iModulons in this category mostly include uncharacterized genes. 4) The iModulons with poorly-matched category (lower left quadrant) have low regulon and iModulon recall. These iModulons contain co-expressed genes that showed statistically significant enrichment levels and appropriate activity profiles [1].

2. **Efflux Pumps**
    a) *czcABC efflux pump:* In our ICA analysis, we identified that the TCS CzcR regulates the CzcR iModulon and is activated in the presence of Zn and Copper (Cu) (**Figure 2b**). Perron *et al*. suggest that CzcS-CzcR TCS is responsible for stimulating the *czcABC* efflux pump, which is involved in heavy metal (Zinc (Zn)/Cobalt (Co)/Cadmium (Cd)) and carbapenem resistance [2]. In our ICA analysis, we also identified that CzcR (RR) regulates the CzcR iModulon (*czcABC* efflux pump) and is activated in the presence of Zn and Copper (Cu) (**Figure 2b** and **c**). It is confirmed by the strong correlation (PCC 0.68) between the *czcR* gene expression and the CzcR iModulon activity (**Figure 2c**). This is further supported by scanning the expression of *czcR* gene with the activity of all the reported 116 iModulons. Additionally, we also find the *czcABC* efflux pump is also specific to Azithromycin antibiotic (**Supplementary Figure 3a**). Thus, czcABC might have some role in Azithromycin-specific resistance in *P. aeruginosa*.

    b) *mexCD-oprJ efflux pump:* As per our analysis, we found that the *esrC* gene regulates the *mexCD-oprJ* efflux (PCC 0.62) (**Figure 2c**). The *esrC* gene is an AraC homolog and a transcriptional regulator controlled by EsrA-EsrB TCS. Furthermore, it was confirmed by checking the correlation of expression *esrC* gene expression with the activity of 116 iModulons (**Supplementary Figure 3b**). Previously, Bruns *et al*. suggest that bioactive compounds of garlic (allyl sulfides) inhibit the EmrD-3 pump-mediated drug efflux in *Vibrio cholerae* [3]. However, in *P. aeruginosa* we find that the presence of antimicrobial agent diallyl disulfide (DADS) and Chloramphenicol led to the overexpression of *mexCD-oprJ* containing iModulon named EsrC.

    c) *mexEF-oprN efflux pump:* MexS-1 iModulon incorporates genes of *mexEF-oprN* efflux pump with negative gene weight (**Figure 2c**). Fetar *et al* show that *mexT* is the activator of *mexEF-oprN* efflux pump [4]. We also found that mexT (RR) controls the *mexEF-oprN* efflux pump containing iModulon (PCC -0.95) (**Figure 2c, Supplementary Figure S4a**). Nies *et al* show that *mexEF-oprN* efflux pumps help in the detoxification of some organic and inorganic substances [5]. The *mexEF-oprN* efflux pump is also known to export the quorum-sensing signals [6]. However, from the conditions, which we used in our study, we find that it is also specific to heavy metals like Zn, Cu, Iron (Fe), as well as organic compound N-acetyl glucosamine (GlcNAc) (**Figure 1c**).

d) *mexGHI-opmD efflux pump:* SoxR iModulon has the genes of *mexGHI-opmD* efflux pump (**Figure 1c, Supplementary Figure S3c**). We identified PA2274 gene that belongs to the OmpR family and has the property of transcriptional regulator using our previously published method [7]. Further, we predicted its role in regulating the *mexGHI-opmD* efflux pump (PCC 0.61) (**Figure 1c**). Aendekerk *et al* showed that *mexGHI-opmD* efflux pump controls antibiotic susceptibility, growth, and virulence [8]. More precisely, we found that *mexGHI-opmD* efflux pumps activate in the presence of *n*-alkanes, biofilms, and beta-lactam (Ceftazidime and Piperacillin) antibiotics.

e) *muxABC-opmB efflux pump:* CpxR iModulon consists of *muxABC-opmB* efflux pump. We identified the PA3205 gene as a potential RR by our previously published method [9] and it's also reported in P2CS database [10]. Further, we checked the relationship between the PA3205 gene and the CpxR iModulon activity and find a strong correlation of 0.84 (**Figure 1c** and **Supplementary Figure S4b**). We find that epigallocatechin gallate and p-anisaldehyde increase the activity of CpxR iModulon, which is also reported by Adewunmi *et al* [11]. Additionally, we also find that *muxABC-opmB* efflux pump is also specific to Zn metal and Ceftazidime (beta-lactam antibiotic) (**Figure 1c**). Thus, we hypothesize that *mucABC-opmB* efflux pump might have some role in beta-lactam resistance in *P. aeruginosa*.

Likewise, we get response regulators *mexZ*, *pprB*, and *brlR* modulating the MexZ, PprB, and CueR iModulons incorporating the *mexXY*, *PA1874-PA1877*, and *mexPQ-opmE efflux pumps* respectively (**Figure 1c** and **Supplementary Figure 2a, 2b, and 4c**). The *mexPQ-opmE* efflux pump containing iModulon shows increased activity in presence of Cu. However, the *PA1874-PA1877* efflux pump containing iModulon shows increased expression in presence of heavy metals (Cu, Zn, Fe) and organic compound GlcNAc. Further, the *mexXY* efflux pump containing iModulon shows less activity in the knock-out condition of *mexZ* gene.

Therefore, the ICA analysis of the *P. aeruginosa* transcriptome would be helpful in the identification of respective TCSs (RR and HK) regulating the efflux pumps. We also identified the exact substrates specific for particular efflux pumps (as detailed in form of heatmap **Figure 2b**)
